# WAVES: a Web Application for Versatile Enhanced bioinformatic Services

**DOI:** 10.1101/276485

**Authors:** Marc Chakiachvili, Sylvain Milanesi, Anne-Muriel Arigon Chifolleau, Vincent Lefort

## Abstract

**Summary:** WAVES is a web application dedicated to bioinformatic tool integration. It provides an efficient way to implement a service for any bioinformatic software. Such services are automatically made available in three ways: web pages, web forms to include in remote websites, and a RESTful web services API to access remotely from applications. In order to fulfill the service’s computational needs, WAVES can perform computation on various resources and environments, such as Galaxy instances.

**Availability and implementation:** WAVES was developed with Django, a Python-based web framework. It was designed as a reusable web application. It is fully portable, as only a Python installation is required to run Django. It is licensed under GNU General Public License. Source code, documentation with examples and demo are available from http://www.atgc-montpellier.fr/waves/.

## 1 Introduction

Any new bioinformatic tool must be made available to its user’s community, essentially to biologists, for whom command line interfaces are often cumbersome. This need is mainly satisfied by implementing a dedicated website. Several solutions, such as Galaxy [3] or Mobyle [4], were developed to ease tool integration within automatically-generated web pages, making them accessible through a generic web user interface. These generic approaches allow the integration of a large variety of bioinformatic tools behind the same interface model. However, each scientific community may also require web applications with specific interfaces adapted to their needs. For instance, phylogeny.fr [1], a well-known pipeline for phylogeny inference, integrates visualization modules dedicated to specific phylogeny data. These specific web applications do not rely on generic solutions which provide too constrained interface frameworks.

Here, we present a versatile service-oriented web application, named WAVES, designed to provide an integrated web-oriented interface for bioinformatic tools, as a facade [2] that conceals the complexity of the underlying computing architecture. The main goal of WAVES is to gather a comprehensive selection of bioinformatic services within a single application programming interface (API). It may integrate tools from different environments and remote resources. In this way, WAVES allows bioinformaticians to integrate tools easily so they can focus on designing high-level user interfaces for community specific web applications. Moreover, WAVES was designed with a focus on facilitating installation and use in production environments.

**Figure 1:**
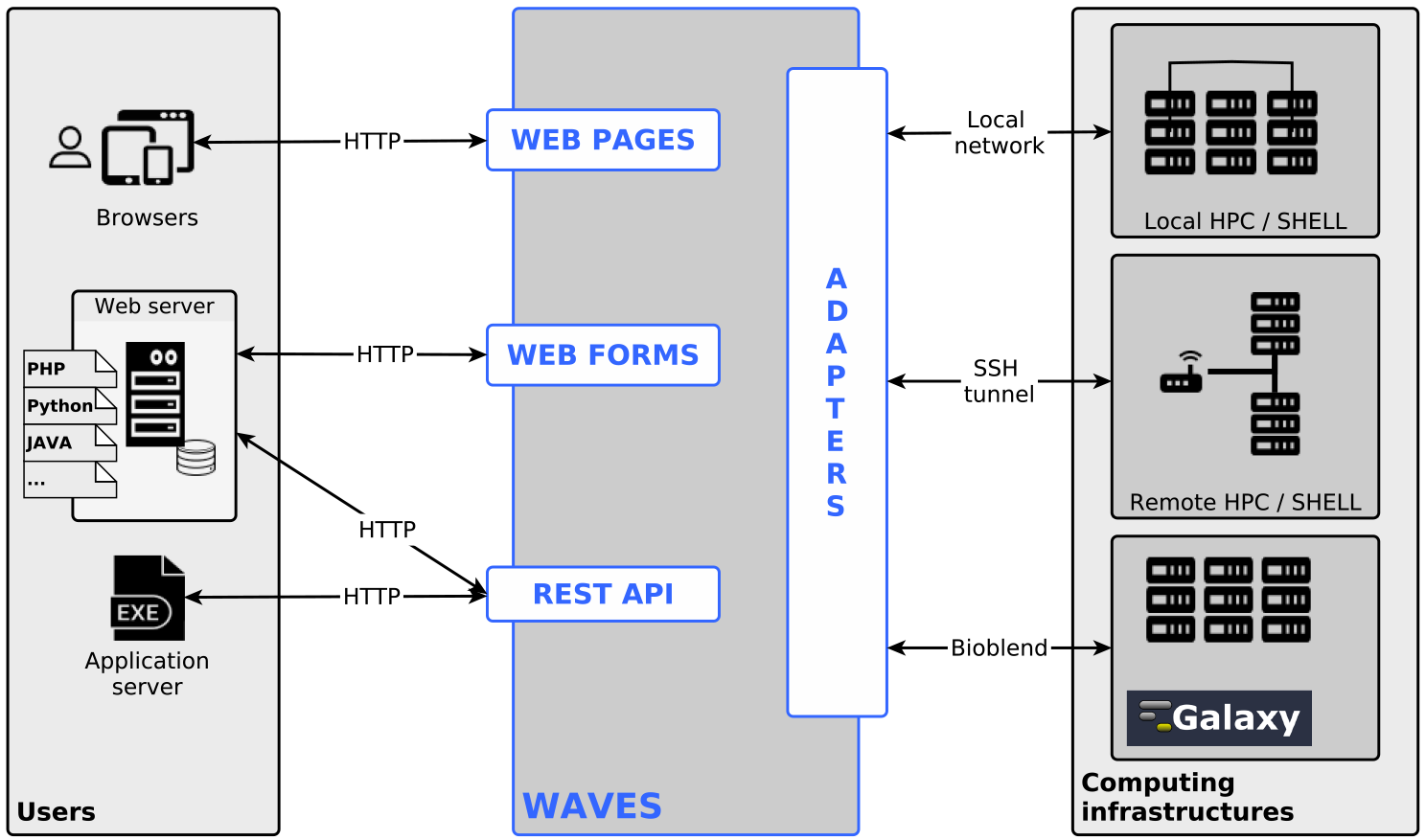
WAVES overview. WAVES provides three different ways for users to access bioinformatic services. It creates “WEB PAGES” for direct user access via web browsers. It also generates “WEB FORMS” to include into remote websites. Lastly, the services are available through “REST API” entries, which are intended to be requested by any remote application (including web servers). WAVES can run the tasks required by services on a variety of local and remote computing infrastructures, including Galaxy instances, using specific “ADAPTERS”.

## 2 Features

WAVES is a web application based on a service-oriented architecture, intended to be used as a service provider platform. These services share a common layout that is unrelated to underlying computational resources. WAVES is a reusable web application designed to run with most web servers, such as Apache and Nginx. It is easily customizable to match the end-user and webmaster needs. Any tool can be integrated into WAVES and configured as a service, provided that tool has well-defined input and output files and exit codes.

There are three different ways to interact with WAVES services: web pages, web forms, and a RESTful API (Figure 1). WAVES automatically creates a web page for each integrated tool. This basic feature is essential for providing end-users with an interface that enables them to run online bioinformatic analyses. In the same manner, it generates web forms to be directly integrated into any website. Lastly, WAVES provides web service entries in its RESTful API, thus generating services suitable for software interoperability. These web services all share the same API structure. Complete details on API architecture and usage are available in Supplementary Material (supplementary Tables 4 and 5).

WAVES is highly versatile, as it is compatible with a variety of computing infrastructures. It runs any locally installed tool. By setting the required credentials, WAVES runs remotely installed tools through a secure network connection. It interoperates with most computing resource management systems, provided that they comply with DRMAA [6]. For interoperability purposes, WAVES interacts with Galaxy, a widely used workflow management system. Thanks to the BioBlend API [5], WAVES lists the tools available in Galaxy instances and offers the ability to import them automatically as new services. WAVES can then run the tools within the Galaxy instance from which it was imported, check computation status, and retrieve results. As WAVES can communicate with several Galaxy instances, tool integration from several locations is straightforward.

WAVES was written in Python using the Django web framework, which complies with web standards. It is fully portable, as it only requires a Python installation and a running web server. It can thus be installed easily into existing web systems. Moreover, it is released as a reusable Django application and may be integrated into any existing Django project.

## 2.1 WAVES as a tool provider

Making tools available on the web is often a thankless and time-consuming task. By taking advantage of WAVES features, this task is greatly simplified, saving time. WAVES can be used as an online service provider. Configuring a tool as a service is achieved by filling out a form in the WAVES administration interface to define the tool settings (supplementary Figure 3). Once set up, a tool is immediately available online through web pages, web forms and via the API. This newly created service can be integrated into remote websites by incorporating web forms. At each service upgrade, corresponding web forms are updated immediately and propagated to remote websites. The same mechanism holds for the API, thus allowing service integration into any remote software. The API can also be used to develop dedicated custom web forms, thus providing high-level user web interfaces.

## 2.2 WAVES as a tool agregator

The existing tool integration solutions provide limited functionnalities to access remote services running on remote computing resources. Mobyle provides remote services integration restricted to services available on remote Mobyle instances, and Galaxy can only run tools installed on its own locally computing resources [3].

Using its interoperability features, WAVES can be used to aggregate remote services behind a common front-end. It can integrate tools easily from remote servers and Galaxy instances. These services rely on remote computing resources maintained by remotely located tool providers. Using WAVES that manner, it is possible to incorporate selected services without the need for a dedicated computing infrastructure, thus saving installation and maintenance costs. Moreover, the tools updates are handled by tool providers, which saves time.

Once freed from locally computing resources maintenance and locally installed bioinformatic tools management, the bioinformaticians in charge of website implementation can focus on developing high-level web interfaces that suit end-user needs. Thus, their expertise in bioinformatics can be entirely devoted to developing dedicated interfaces.

